# Comparative RT-qPCR and qPCR reveals early infection, low-titer infection, and relative cell activity of the HLB bacterium, *Candidatus* Liberibacter asiaticus

**DOI:** 10.1101/2024.11.05.622139

**Authors:** Rachel Patterson, Desen Zheng, Weiqi Luo, Zhanao Deng, Clive Bock, Ruhui Li, Yongping Duan

**Affiliations:** USDA-ARS, US Horticultural Research Laboratory, Fort Pierce, Florida, 34945, USA; Center for Integrated Pest Management, North Carolina State University, Raleigh, NC 27695, USA; Gulf Coast Research and Education Center, University of Florida, Wimauma, FL 33598, USA; USDA-ARS, National Germplasm Resources Laboratory, Beltsville, MD 20705, USA

**Keywords:** Citrus Huanglongbing, *Ca.* Liberibacter asiaticus, RT-qPCR, qPCR, early detection, Low-titer infection, and relative cell activity

## Abstract

*Candidatus* Liberibacter asiaticus (Las) is one of the causal agents of citrus huanglongbing (HLB) epidemics worldwide. Due to its fastidious nature, intracellular and systemic infection, detecting Las at early and/or low-titer infection, as well as differentiating between live or dead cells in the host psyllids and citrus plants is critical for effective HLB management. To achieve both sensitive Las detection and differentiation, we employed one-step reverse transcription-quantitative PCR (RT-qPCR) using total nucleic acids as template. This method allows use of both Las 16S rRNA and rDNA as template in the same reaction and increases detection sensitivity by up to 1000-fold in comparison with quantitative PCR (qPCR). The increased sensitivity significantly reduces false negative detection and detects the otherwise undetectable low-titer Las infections. Furthermore, the greater the abundance of 16S rRNA present in the samples, the bigger the Ct gap obtained between RT-qPCR and qPCR results. The numerical Ct gap can be used to deduce relative Las cellular activity and indirectly infer whether cells are alive or dead. In addition, this comparative detection method also can be used to select inoculum and monitor relative cell activity during *in vitro* Las culture and evaluate the effectiveness of antimicrobial treatments against Las bacteria.

## Introduction

Citrus Huanglongbing (HLB) is a devastating disease of *Citrus* spp., causing billions of dollars in losses annually worldwide (Li et al. 2020). HLB is caused by any of three fastidious α-proteobacteria, namely *Candidatus* Liberibacter asiaticus (Las), *Ca.* Liberibacter africanus, and *Ca.* Liberibacter americanus which are vectored by citrus psyllids. Among them, Las is the most destructive and widespread globally. With no cure or known resistant commercial varieties (Das et al. 2019), the search for a rapid and high-throughput early detection method of very low titers to mitigate disease impact and target infected trees for quarantine or removal, as well as to support the distribution of truly Las-free nursery stock, is of the utmost importance (Bao et al. 2020).

However, the fastidious nature of Las and limitations in current diagnostic methodology hinder detection of early and/or low-titer infections. The incubation period for low-titer infections may be longer (Pandey and Wang 2019), sometimes taking years to show symptoms whilst still providing an inoculum source to vectors (Lee et al. 2015). Provided that low-titer infections are involved in the development of disease and spread of the bacteria and that Las colonizes trees unevenly (Kunta et al. 2014a; Tatineni et al. 2008), there is uncertainty in representative sampling and the possibility of false negatives in trees that would otherwise be rogued or quarantined when the titer is below the limit of detection (Morgan et al. 2012; Selvaraj et al. 2018; Wheatley et al. 2021). Although other techniques have been proposed (Arredondo Valdés et al. 2016; Pandey and Wang 2019; Weng et al. 2022), quantitative PCR (qPCR) and its variants remain the most viable option for the reliable detection of Las given their availability, accuracy, and adoptability (Das et al. 2019). The qPCR-based method published by Li et al. in 2006 has been used as the standard method for the detection of Las for more than 18 years. However, deploying new technological advances may improve its diagnostic applications (Das et al. 2019; Osman et al. 2023; Selvaraj et al. 2018).

RT-qPCR is often used for gene expression studies and the detection of RNA viruses (Rocha et al. 2020) but it is also useful for enhanced sensitivity in detecting microbes (Guirou et al. 2020; Kriegova et al. 2020; Pitkänen et al. 2013), distinguishing viable bacteria, and determining cellular activity (Freitas et al. 2022; Miao et al. 2018; Pitkänen et al. 2013; Váradi et al. 2017). While conventionally paired with RNA extracts (Kim and Wang 2009), the use of RT-qPCR with total nucleic acid extracts (TNA) shows promise for enhanced sensitivity in detection (Guirou et al. 2020; Minguzzi et al. 2016) and correlation to relative cellular activity (Pitkänen et al. 2013). Due to the known sensitivity of RT-qPCR, it has been used to declaratively report the absence of bacteria in samples (Kuperman et al. 2020).

The ability to distinguish live and dead cells is crucial to ongoing research on HLB. Antimicrobial testing of the fastidious Las bacteria relies on differentiation between viable and nonviable cells, which can be approximated by measurement of relative cellular activity. Traditional qPCR, while sensitive, is not able to distinguish between DNA from active cells and that of dead or otherwise inactive cells (Cangelosi and Meschke 2014). In the absence of a method for pure culture, previous attempts to make this distinction have involved viability PCR (vPCR) using an additional ethidium monoazide (Trivedi et al. 2009) or propidium monoazide step (Hu et al. 2013; Louzada et al. 2022; Vazquez 2015). One-step RT-qPCR, in contrast, reverse-transcribes RNA from a total nucleic acid extraction into cDNA that is then available for fluorescent tagging in the qPCR step along with the original double-stranded genomic DNA fragments (Adams 2020). Due to RNA’s instability and shortened half-life relative to DNA, inactive cells will have degraded RNA not amenable to reverse-transcription. The addition of RNA from active cells may lower the Ct in RT-qPCR, giving a relative measure of cellular activity and active bacterial titer (Miao et al. 2018) and thus much more sensitive detection.

In this study, we applied RT-qPCR and compared it with qPCR using TNA from infected Asian citrus psyllids (ACP) and citrus plants. Sensitivity of the one-step RT-qPCR was much higher, which allows the detection of otherwise undetectable samples from early infection or low-titer infection of Las. By comparing RT-qPCR with qPCR Ct results, the relative cell activities of Las bacteria can also be deduced from the difference between Ct gaps.

### Materials and Methods

### Citrus and psyllid samples

Plants were maintained in the USHRL insect-proof greenhouse in Fort Pierce, FL, and continuously inoculated with Las+ psyllids and/or graft-based inoculation with side grafting (Zhang et al. 2012) to maintain Las populations *in planta*. Leaves were sampled by collecting 3-5 leaves from a single tree with >20 replicates. Citrus samples were divided into specific collections of symptom status (asymptomatic, blotchy mottle, and yellow shoot) that were identified and collected by highly experienced HLB researchers, and into a category of “general” collections replicating in-field samples sent for diagnostics that may combine asymptomatic/symptomatic and Las+/Las-samples into one. Additional sets of asymptomatic leaf samples were collected separately from individual branches on greenhouse grown Valencia and navel sweet orange trees for detection of early infections at 49-, 120-, and 187-days post-graft inoculation (dpi) with Las-infected bud-sticks. Three leaves from each of 75 Valencia branches from 15 individual trees and 25 navel orange branches from 5 individual trees was collected and used in the study for early detection.

Psyllids were reared in the ARS-USHRL Subtropical Insects and Horticulture Research Laboratory (Hall and Hentz 2019), or directly collected from trees displaying HLB symptoms at the ARS-USHRL Picos Research Farm using an insect aspirator.

Cultures were prepared as previously published (Zheng et al. 2024), and sampled by collecting a 500 μL subsample of the liquid culture via pipette.

### Nucleic acid extraction

TNA was extracted from leaf and psyllid samples using a modified cetyltrimethylammonium bromide (CTAB)/column TNA purification method with a water eluent. The CTAB method involved initially mincing 0.1 g tissue into <1 mm^3^ pieces before adding 800 μL of extraction buffer (3% CTAB, 100 mM Tris-HCl, 20 mM EDTA, 1.4M sodium chloride, 5% polyvinylpyrrolidone, pH 8.0) and 80 μL 2% sodium dodecyl sulfate (SDS), disrupting cells in a Fast Prep-24 (MP Biomedicals, Solon, OH, USA) for 2 minutes at 6.5 m/s, and incubating at 65°C for 20 minutes. Samples were treated with 400 μL potassium acetate, vortexed, and incubated on ice for 5 minutes. The lysate was clarified by centrifuging at maximum speed for 5 minutes, decanting the supernatant into a new microcentrifuge tube, and repeating the centrifugation. The supernatant from the second centrifugation was removed, combined with 900 μL ice cold 95% ethanol, and transferred to a UPrep Universal Spin Filter Column (Genesee Scientific, El Cajon, CA) followed by two washes of the spin filter column with 70% ethanol. The filter was dried by centrifuging at 10,000 rpm for 2 minutes. The dried filter was incubated for 5 minutes with 50 μL nuclease-free water to elute TNA before centrifuging at 10,000 rpm for 2 minutes. Individual psyllids were extracted using the modified CTAB method described above with a 55°C incubation and 30 μL of nuclease-free water eluent. Total nucleic acids were extracted from cultures by a simple osmotic shock method (Masson et al. 2018).

### Quantitative PCR (qPCR)

qPCR reactions were performed in an ABI 7500 Fast Real-Time PCR System (Applied Biosystems, Foster City, CA). The primers and probes listed in Supplementary Table 1 were used to detect Las using qPCR. The Li Las primers/probe were used at the concentrations initially published (Li et al. 2006), and combined in 15 μL reactions with 2 μL TNA template and TaqMan^™^ Fast Universal PCR Master Mix (Thermo Fisher Scientific, Waltham, MA) according to the manufacturer’s instructions. Cycling conditions were 95°C for 30 seconds followed by 40 repeats of 95°C for 3 seconds and 60°C for 30 seconds. The Luna^®^ Universal One-Step RT-qPCR Kit (New England BioLabs, Ipswich, MA) was also used for qPCR without the reverse transcription step in cycling with the same concentration of the Li Las primers/probe set. Cycling conditions were optimized for additional primer/probe sets after an initial annealing phase of 95°C for 1 minute; 40 cycles of 95°C for 5 seconds and 60°C for 30 seconds for the RNR primers/probe (Zheng et al. 2016) (Supplementary Table 1), and 40 cycles of 95°C for 3 seconds and 62°C for 30 seconds for the LJ-900 primers/probe (Morgan et al. 2012) (Supplementary Table 1). Ct values were calculated by the Applied Biosystems 7500 Real-Time PCR Software version 2.3 with a standard threshold of 0.02 ΔRn. Ct values below 24 were considered to be high-titer, between 24 and 28 as medium-titer, and between 28 and 36.9 as low-titer. The amplification efficiency was determined using serial dilutions of high-, medium-, and low-titer leaf samples and psyllid samples in ten-fold dilutions of a 100 ng sample with either water or uninfected citrus TNA as the diluent.

### Quantitative reverse transcription PCR (RT-qPCR)

RT-qPCR reactions were prepared and amplified using the Luna^®^ Universal Probe One-Step RT-qPCR Kit (New England BioLabs, Ipswich, MA) according to the manufacturer’s protocol, with a reduction from 10 seconds to 5 seconds in the denaturation step of the Li primer protocol. RNR and LJ-900 were cycled as in the qPCR methods described above, with a preceding cDNA synthesis step of 55°C for 10 minutes. Ct values were calculated by the Applied Biosystems 7500 Real-Time PCR Software version 2.3 with a standard threshold of 0.02 ΔRn. Samples were considered high-titer if the Ct value was 24 or less, medium-titer if the C_t_ was between 24 and 28, and low-titer between 28 and 36.9. The amplification efficiency was evaluated as for qPCR.

### Comparison of three sets of primers and probes for sensitivity

The Li primers and probes have been widely adapted due to their sensitivity and specificity to the 16S rDNA subunit of the three HLB-causing *Ca.* Liberibacter species (Li et al. 2006). However, other primers and probes have been developed and tested for specificity and sensitivity to Las. Among these are the LJ-900 primers and probe targeting the *LasA_I_*/*LasA_II_* prophage region of the Las genome (Morgan et al. 2012) and the RNR primers and probe targeting the Las *nrdB* gene, the β-subunit of ribonucleotide reductase (Zheng et al. 2016). RT-qPCR-based diagnostic assays may operate with a number of primer sets, so to determine which of the more extensively verified primer and probe sets would afford the most assay sensitivity we tested all three in side-by-side comparisons involving different host tissues. Samples were selected to cover a range of Las titers, as tested in Li RT-qPCR, to avoid true negatives while verifying the primer activity across multiple titers. We were most interested in testing the hypothesis that 16S rRNA/DNA targets (Li) are more sensitive to differences in rRNA production than DNA-specific targets, given the possibility that rRNA could be contributing to the enhanced sensitivity. In this experiment, individual citrus and periwinkle leaves were tested in side-by-side reactions with each of the three primer and probe sets in both RT-qPCR and qPCR.

### Calculation of Ct gaps

RT-qPCR and qPCR values were compared to produce a value, the “Ct gap,” describing the reduction in Ct values from qPCR to RT-qPCR analysis of the same sample. Ct gaps were calculated using the equation *Ct*_*gap*_ = *Ct*_*qPCR*_ − *Ct*_*RT*−*qPCR*_. It was estimated that each increment of one in the Ct gap corresponded to a three-fold change in the bacterial titer.

### Statistical analysis

Statistical processing and presentation of the data generated by the project was performed in MS Excel (Microsoft, Inc., Redmond, WA) and R version 4.2.2 (R Core Team, 2024). Prior to statistical analysis, preliminary data exploration showed that no further transformation was required to satisfy the normality and homogeneity of variance assumptions of the proposed statistical methods. All qPCR Ct results displaying the “Undetermined” value with any of the primer pairs (Li 16S, LJ-900, RNR) were arbitrarily assigned the value of 40, indicating zero detection after 40 cycles. Las levels above a Ct of 36.3 were considered negative. Bar charts and boxplots across multiple trials were employed to visualize the comparative performance of qPCR and RT-qPCR, using samples from citrus, periwinkle, or ACP at different time points post-inoculation. Bar charts illustrated the overall detection percentages, while boxplots displayed the Ct values and their variability within each trial, providing a comprehensive comparison of both techniques over time. The effect of host types (citrus, periwinkle and ACP), inoculation source, primer sets, and days after inoculation on Las detection accuracy were analyzed by analysis of variance (ANOVA), followed by a post hoc test using Tukey’s honestly significant difference (HSD) tests (α = 0.05) to compare means.

## Results

### RT-qPCR dramatically increases detection sensitivity, especially for low-titer infection

Low-titer infections are a relatively understudied phenomenon as the techniques available to diagnose them are sparsely published. To compare detection capabilities between the original qPCR method and the new RT-qPCR method, initial tests were performed to determine the sensitivity of each assay as illustrated by the lowest amount of TNA template using samples of categorically different titers. Initially, samples were tested by qPCR and the titer classification was determined as either high, medium, or low. One high-titer, one medium-titer, and one low-titer leaf TNA sample each were selected for further testing as was one high-titer psyllid.

Samples were serially diluted with nuclease-free water from 100 ng to 0.1 pg of TNA per well, and these dilutions were used to determine the least amount of TNA detectable by the method. TNA samples with a higher proportion of Las TNA in the sample, regardless of host, remained detectable to 1 pg TNA total using RT-qPCR, while samples with a low proportion of Las TNA were detectable to 1 ng. Samples with a moderate titer/proportion of Las TNA were detected with as little as 100 pg TNA template using the RT-qPCR reaction. When using qPCR, detection was 10-100 times less sensitive, depending on the proportion of Las TNA in the sample. With lower proportions of Las TNA to host TNA, detection was limited to 100 pg-1 ng of TNA (data not shown). Additionally, the improvement in sensitivity RT-qPCR with TNA was very precise. Regardless of the initial host titer (**Error! Reference source not found.**), gaps were consistently an average of ∼4.97 cycles in citrus and ∼7 cycles in psyllids.

Dilution with water is useful for determination of general sensitivity but, due to the obligate biotrophy of the phloem-limited Las bacteria, any nucleic extract of Las will be part of a mixed population of nucleic acids from different organisms with host nucleic acids the most abundant within the sample. To account for the high host-background in samples, we additionally produced a dilution series of a Las-high-titer citrus leaf TNA extracts diluted with uninfected citrus TNA. This testing confirmed that, even with a high background of host TNA, as little as 1 pg of high-titer TNA template is sufficient for detection of Las.

Given the increased sensitivity of RT-qPCR compared to qPCR, we performed further testing to determine whether detection of naturally occurring samples would be improved by RT-qPCR as well. TNA samples extracted from psyllids, periwinkle, or citrus were analyzed using both methods and the number of samples detected compared between the methods. While no samples were determined to be Las positive with qPCR that were determined Las negative with RT-qPCR, the reverse was true for 18.40% of all samples (Figure 2). This indicates that the superior detection capability of RT-qPCR may extend to field samples more generally. In terms of host, detection was improved by RT-qPCR in psyllid samples (16.16% higher detection rate) and citrus samples (15.69% higher detection rate), and vastly improved in periwinkle samples (40.0% higher detection rate).

**Figure 1.**
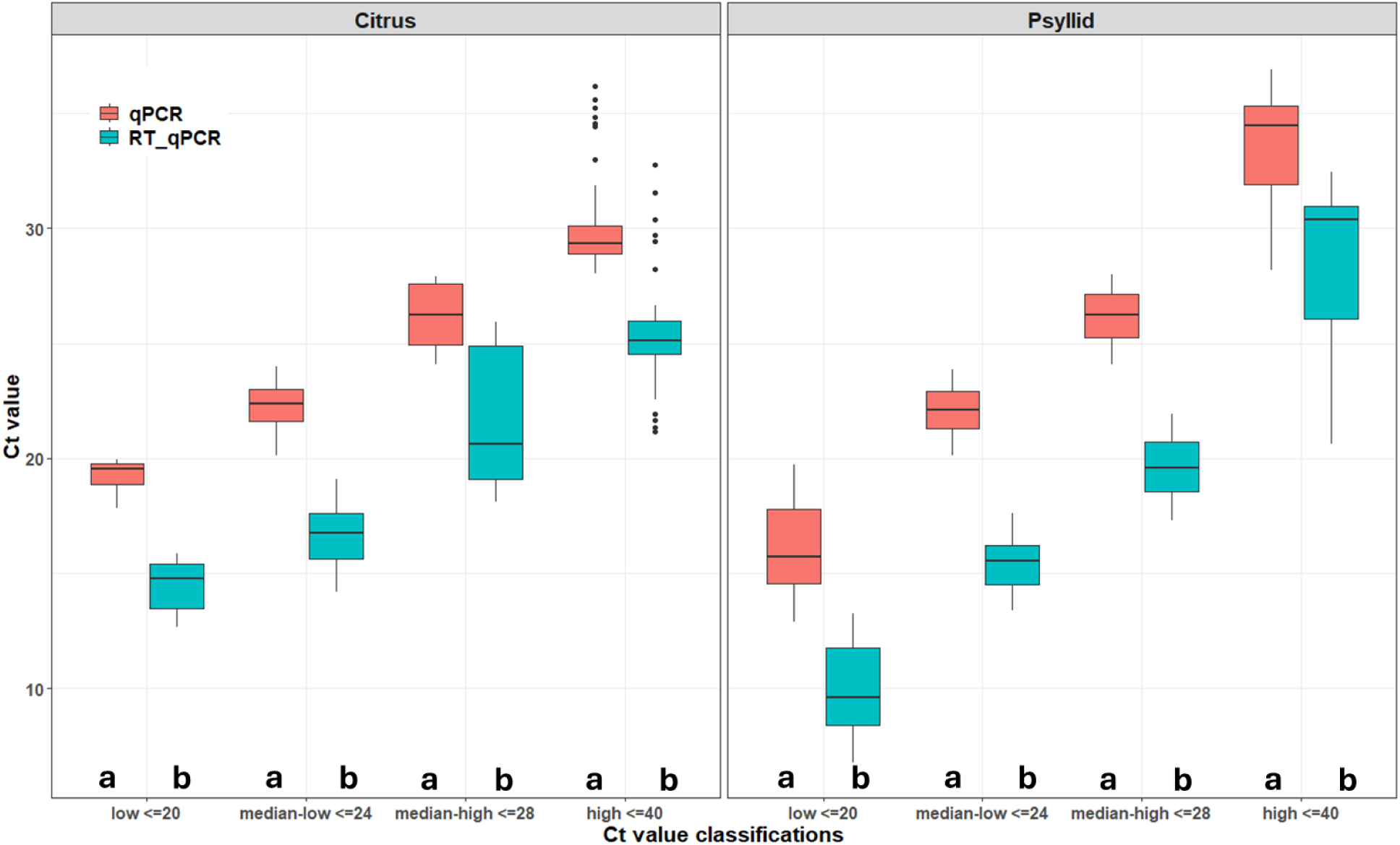
Ct value comparisons with different Las titers detected by qPCR and RT-qPCR. A) Comparison between qPCR and RT-qPCR for different citrus samples with known Las titers; B) the same comparison in psyllid samples. A significant difference between qRCR and RT-qPCR is indicated by different letters.

**Figure 2.**
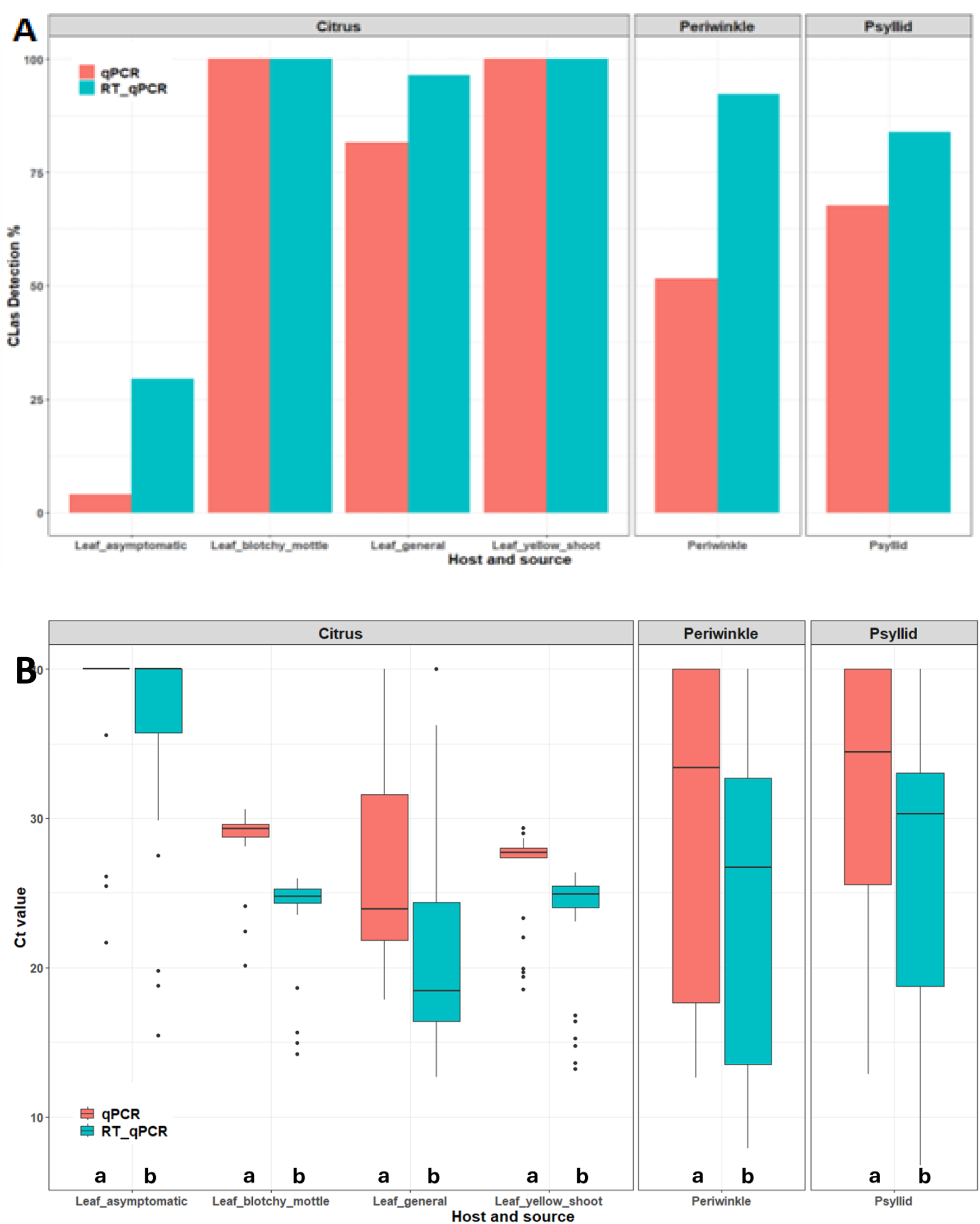
Comparative detection of Las in plant and psyllid samples. A) Detection percentage of Las in plant and psyllid samples; B) the Ct values obtained by either the one-step RT-qPCR assay (blue column) or standard qPCR assay (red column). The samples were collected from known Las-positive sources with different classifications. A significant difference between qRCR and RT-qPCR is indicated by different letters.

Analysis of sample sources indicated significant improvements in detection among specific tissues in citrus and among collection sources of psyllids. General collections, as expected, had more variable titer detections though in all samples RT-qPCR was more sensitive to lower relative titers of Las. For these general collections, qPCR resulted in false negatives in 15.45% of cases. Psyllids generally carry a high titer of Las in HLB endemic Florida groves (Hall 2018). Although the number of detections was equivalent in natural collections, the means of the same samples in qPCR were approximately **7** cycles greater than those of RT-qPCR, indicating that the sensitivity of the detections was greatly improved by RT-qPCR.

### The sensitivity of RT-qPCR depends on the target gene

In comparing the three most sensitive PCR-based detection systems side by side with the same reaction conditions and program settings, the LJ-900 and RNR systems showed slightly increased sensitivity in qPCR using the DNA as template as reported by Morgan et al. (2012) and Zheng et al. (2016). When using RT-qPCR with TNA as template, the Li system was the only system that dramatically increased the detection sensitivity, as demonstrated by the Ct values decreasing by >8.0 cycles (Figure 3). Although high-titer samples were detected by all three set of primers and probes, low-titer leaves were more clearly detected with RT-qPCR using the Li primers and probe. In addition to the number of samples detected, the Ct values for the samples were significantly lower with RT-qPCR when using the 16S rRNA/rDNA as template than either the RNR or LJ-900 primers and probes (**Error! Reference source not found.**). These results indicate that the 16S rRNA accumulated significantly more in the reactions than the transcripts of the *LasAI* and

**Figure 3.**
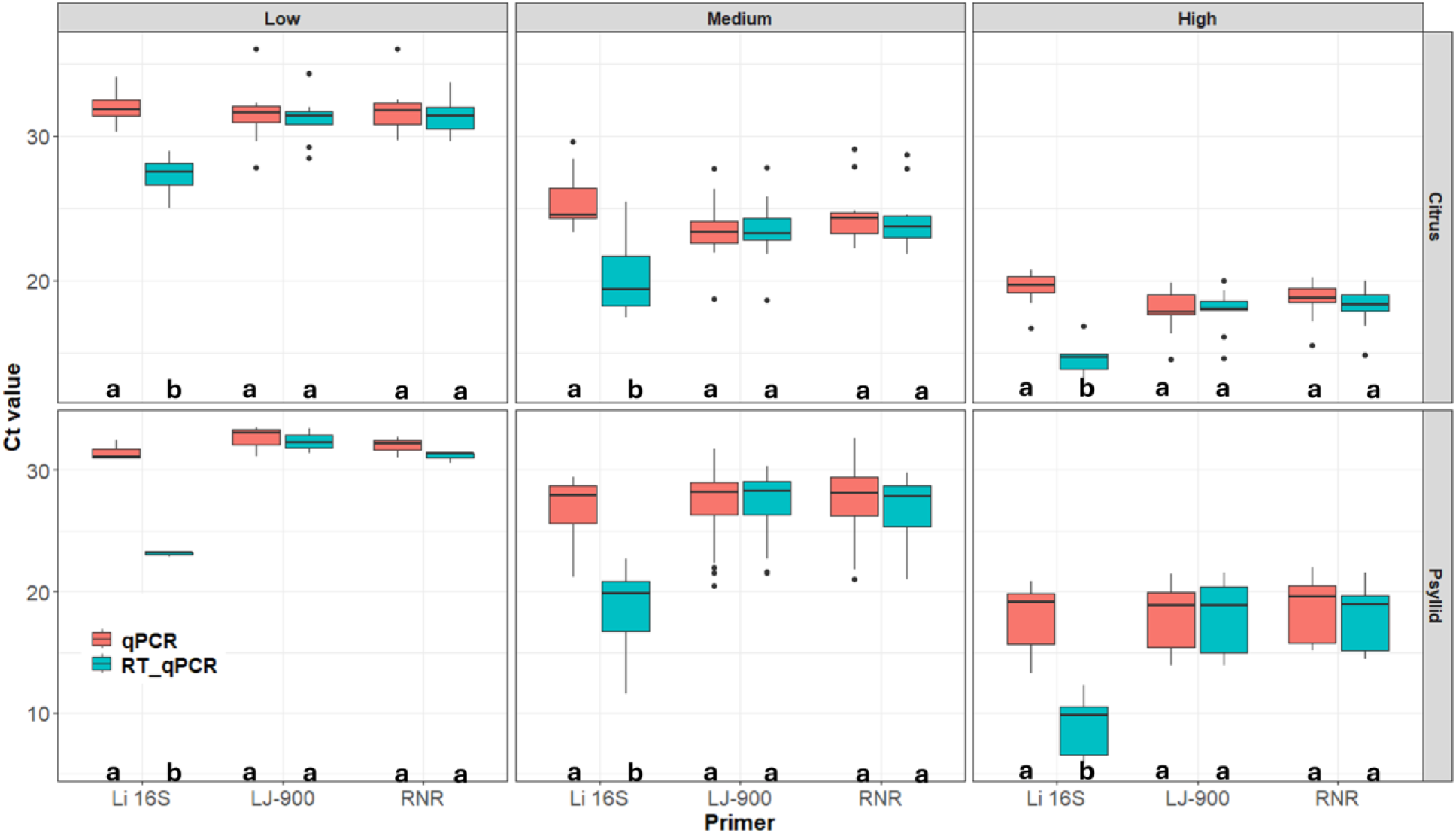
Sensitivity comparison among three different sets of primers and probes. A) Li primers and probe display much more sensitivity across low, medium and high titers of Las when RT-qPCR was used. B) Both LJ-900 and RNR primers and probes show slightly more sensitive when qPCR was used, but no difference between qPCR and RT-qPCR for both. A significant difference between qRCR and RT-qPCR is indicated by different letters.

*LasA_II_* transporter genes and the β-subunit of ribonucleotide reductase gene. Therefore, the 16S rDNA/rRNA target is an optimal target for the RT-qPCR assay.

### Low-titer infection is either transitory or persistent in planta

Low-titer infection can be “transitory,” meaning that early infection will develop to high titer infection with typical HLB symptoms (Figure 5Bc). Given the heightened sensitivity of the RT-qPCR assay with the widely used Li primers, the possibility of detecting Las in plants prior to the appearance of symptoms was chosen for exploration. Of the 100 asymptomatic leaf samples taken from plants 49 days-post-inoculation with Las (dpi), only 3 were detectable using standard qPCR with an average Ct of 29.05 ±2.67. However, RT-qPCR detected Las in 29 of samples at an average Ct of 32.62 ±0.79 in the same samples. At 77 dpi, 89% of the 55 asymptomatic leaf samples were detected in RT-qPCR with an average Ct of 33.75 ±0.20 (Figure 4).

**Figure 4.**
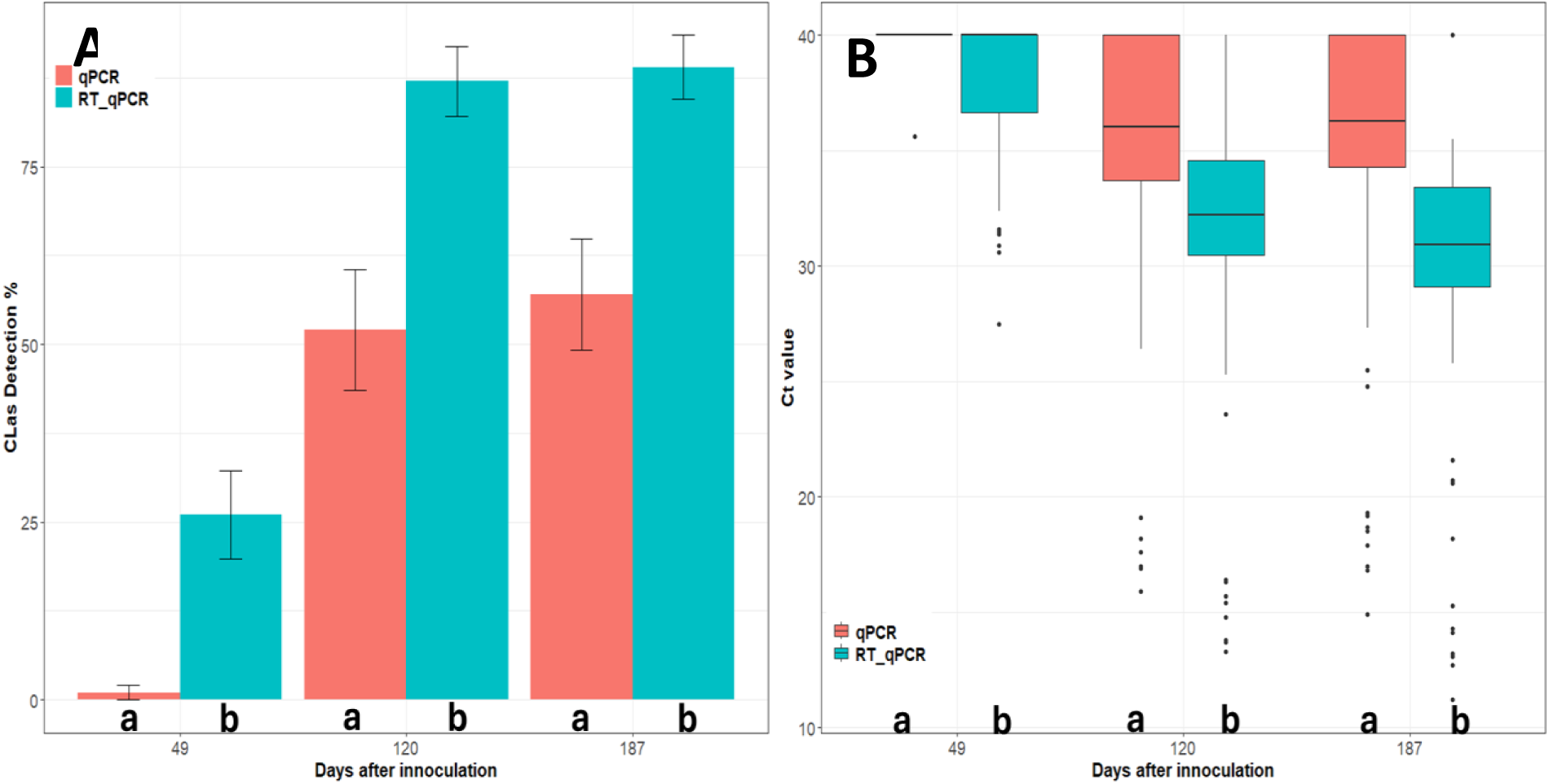
Early detection of Las during its progression in sweet orange citrus plants. A, Las detection percentage and B, Ct values for citrus plants at 49, 120, and 187 days after inoculation using RT-qPCR and standard qPCR assays. A significant difference between qRCR and RT-qPCR is indicated by different letters.

Another mode of low-titer infection is “persistent”. Persistent low-titer infections were prominent in Las-infected grapefruit plants that were inoculated with cultured Las bacteria (Zheng et al., 2024). When budwood taken from 4-year-old low-titer Las-infected plants was used to graft-inoculate 27 healthy Duncan seedlings, after 6 months 12 out of 27 seedlings tested Las+ using RT-qPCR with a range of Ct values from 28.6 to 35.9. Only one of those seedlings tested Las+ using qPCR with a Ct value of 36.2. The same phenomenon was observed in two different sets of grapefruit plants when the plants were treated with penicillin and streptomycin as described by Zhang et al. (2012), propagated, and monitored for more than 6 months and two

These Las-infected plants displayed markedly smaller fully expanded leaves, and/or growth retardation within either a branch or the whole tree (Figure 5Ab, Bb, C and D) as compared to the routinely larger healthy grapefruit leaves (Figure 5 Aa and Ba). The leaves of a grapefruit tree infected with Las at a qPCR-determined average Ct of 34.85 the small leaf branch (Figure 5, C and D) demonstrated Cts ranging from 26.8 to 35.7 in RT-qPCR in the group of 14 out of 24 plants from 6-month group and 3 out of 12 plants from the >2-years-old group. The diagnostic symptoms of “persistent” low-titer Las infections were often small leaves, mild stiffer elastic tissue, and either a lack of or mild chlorosis.

**Figure 5.**
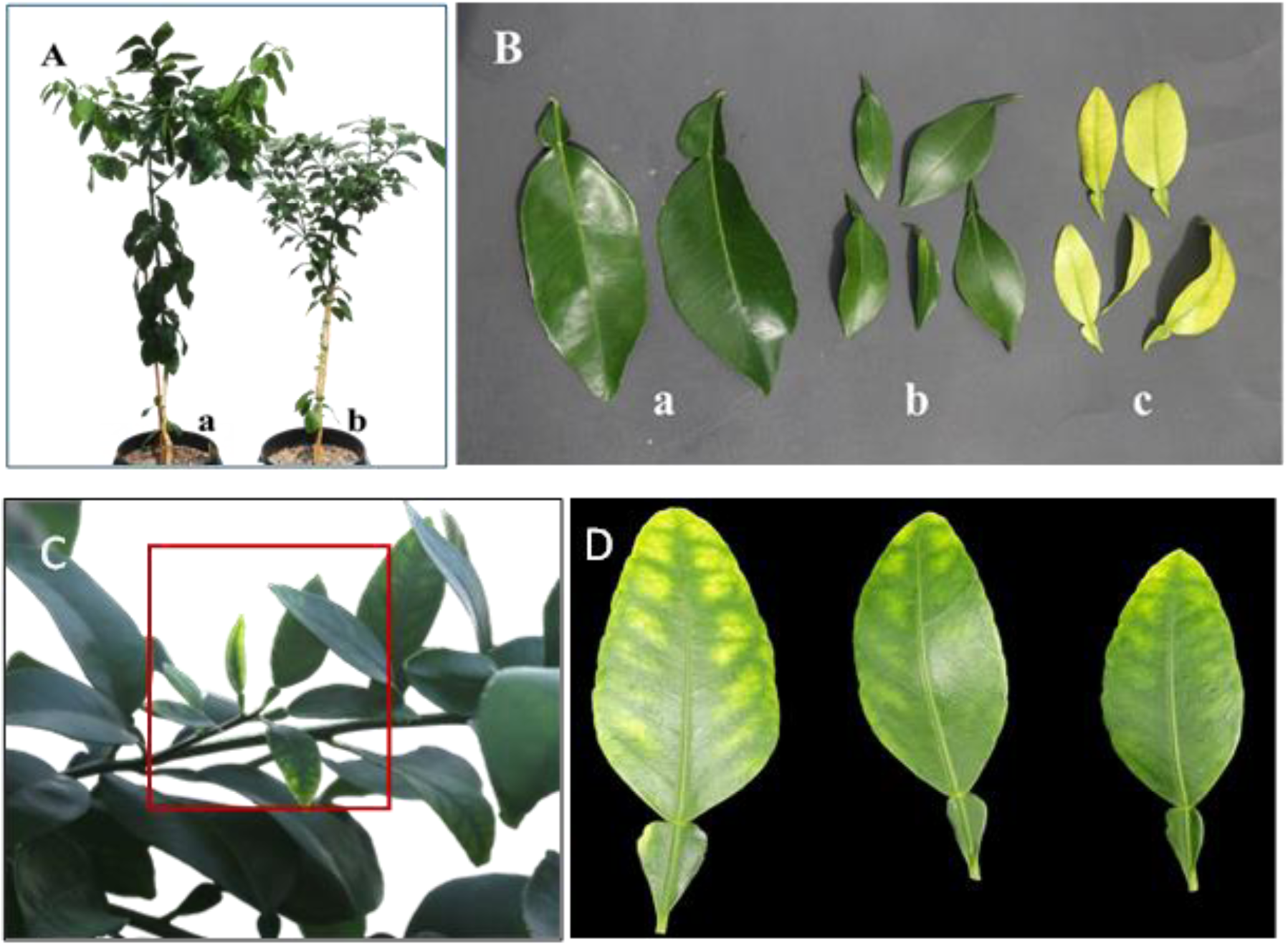
Small leaf symptoms associated with different titers of Las infection. A, Small leaves and growth retardation with low titer infection of Las (Ab) and control Ruby Red grapefruit (Aa) at 2 years old; B, typical HLB symptom with small, yellow leaves and high titer of Las infection (Bc), atypical HLB symptom with small leaves only and low titer of Las infection (Bb) and control healthy grapefruit leaves (Ba). Smal leaf with low titer infection of Las. C) a brachy of over two-year-old grapefruit tree shows small leaves of HLB symptom. D) Close-up pictures of the small leaves

### The Ct value gaps mirror expected relative cellular activity of Las

We initially observed an increase in detection sensitivity with RT-qPCR although, notably, the increased sensitivity was not consistent for all samples tested. RT-qPCR is likely more sensitive with 16S primers due to the relative abundance of rRNA, which is usually an indicator of cellular activity. Considering the likely correlation between RT-qPCR and cellular activity, and the standard use of qPCR to identify 16S rDNA titers, we developed a hypothesis that the gap between the Ct values for the two methods is likely a relative measure of 16S rRNA productivity and therefore of Las cellular activity within a sample. Investigation of the gaps between qPCR and RT-qPCR Ct values for the same samples demonstrated that the two values are on average significantly different between hosts. In this study, periwinkle samples harbored higher titers of Las on average, with psyllid titers being more moderate, and citrus samples generally displaying the lowest titers. However, the gaps between the Ct values did not follow this trend. The average gap between RT-qPCR and qPCR results for periwinkle was 4.78 ±0.90 cycles, for citrus 4.97 ±1.44 cycles, and 5.74 ±1.25 cycles for psyllids (Figure 2).

To reveal the association of symptoms with Las cell activity, asymptomatic leaves from early infections and symptomatic leaves presenting either yellow shoot or blotchy mottle were collected both in the greenhouse and the field by researchers experienced with HLB diagnosis, and the average gaps in Ct values were analyzed. The gaps between qPCR and RT-qPCR Ct values were, on average, 5.80 ±1.04 for those asymptomatic leaves with early infections advanced enough to be detected in both PCR assays. For symptomatic leaves, the average gap was significantly lower at 3.93 ±1.32. This indicated that cellular activity may be lower in established, symptomatic infections due to phloem damage and Las cell death. Blotchy mottle tends to present on older leaves, while yellow shoot is more a malady of new flush. Symptomatic leaves were analyzed separately based on the symptom displayed, and the gaps in Ct values were higher in blotchy mottle-afflicted leaves (4.68 ±0.93) than leaves displaying yellow shoot (3.31 ±1.38) but this was not statistically significant.

While symptom determinations were made by seasoned HLB researchers, more general collections were made as well to simulate collections from both individuals less experienced with identifying HLB symptoms and individuals focusing on whole-tree sampling. The spread in the Ct gaps of leaves from general collections ranged from 2.06 to 8.69, which demonstrated far greater variability than HLB researcher collections of specific pre-symptomatic/symptomatic tissues (Figure 2). Thus, Ct gaps in mixed collections of infected and uninfected leaves will not accurately approximate the relative cellular activity. Whole-tree sampling may not be amenable to this approach of analysis, which is consistent with observations of high variability between branches and leaves of infected trees.

To further investigate the relationship between Ct value gaps and potential bacterial activity, *in vitro* cultures of Las were subsampled and tested over the time course in both qPCR and RT-qPCR side-by-side (Figure 6). For ACP Las culture, the difference in Ct values between qPCR and RT-qPCR remained relatively stable over the 21-day observation period, which was correlated with Las bacterial growth. For citrus Las culture, along with the increase of Ct values, the difference between qPCR and RT-qPCR Ct values reduced much faster in Las-infected citrus samples as compared to cultures derived from Las-infected ACP, indicating rapid breakdown of RNA and lower to no relative cell activity, and potentially the death of Las bacterial cells. After 14 days, the plant-derived culture had no observed gap between Ct values. In addition, the effectiveness of antibiotics against Las were evaluated in our *in vitro* culture system using the Ct value comparison between RT-qPCR and qPCR. Many antibiotics such penicillin and streptomycin that were effective in *in planta* assays were not effective in killing Las *in vitro* (Zheng et al., 2024). Only tetracycline and oxytetracycline were effective in suppressing Las growth, with reductions of Ct gaps to <1 cycle after 14 days (Figure 6, C and D).

**Figure 6.**
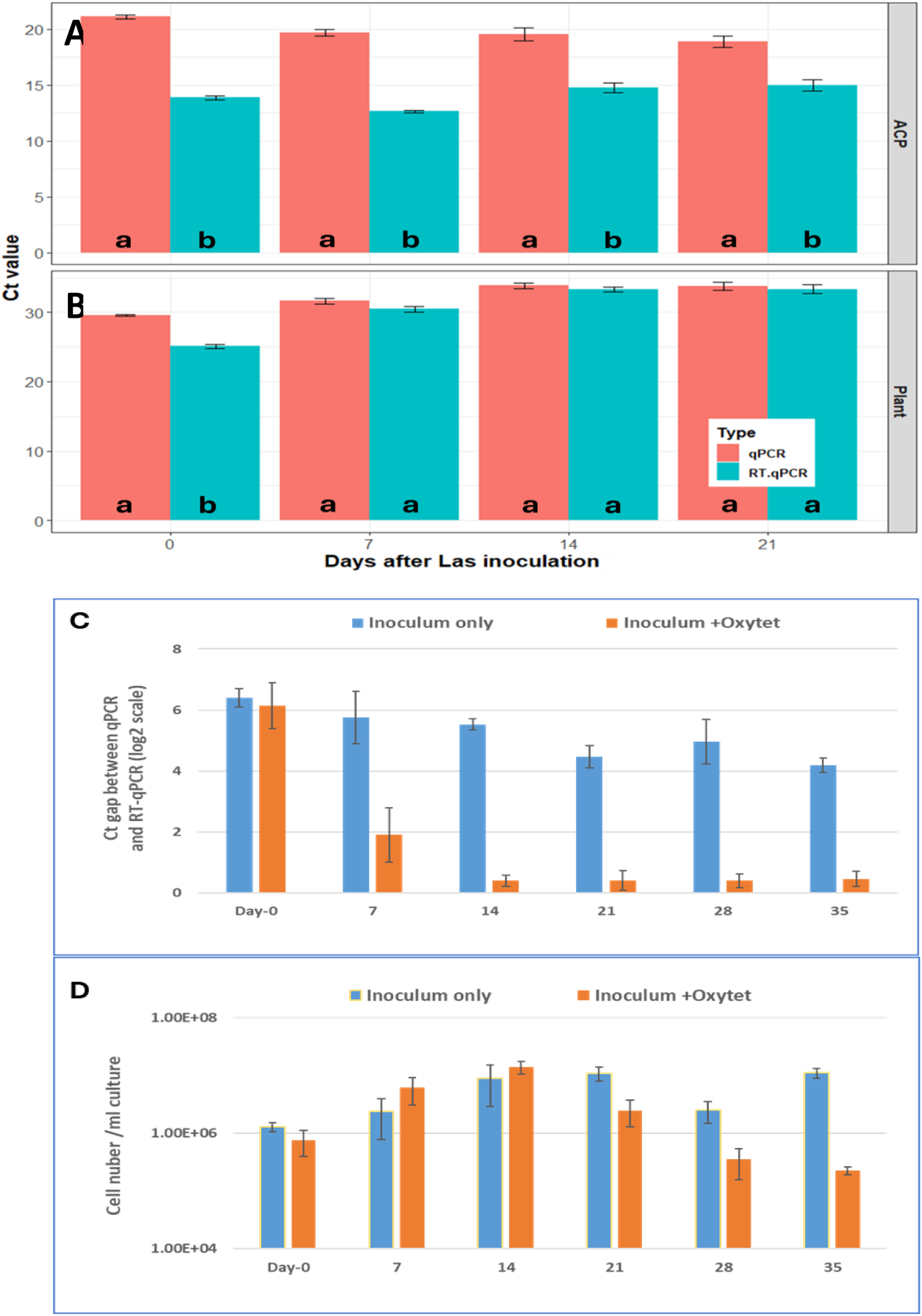
Las growth in vitro culture and their relative cell activity and inhibition effects of oxytetracycline (Oxytet) on Las relative cell activities and growth. Las Ct values detected at 0, 7, 14, and 21days after culture in vitro for ACP-Las samples (A) and citrus-Las samples (B), respectively, using qPCR and RT-qPCR assays. (C), Reduction of Las relative cell activities measured by the gabs between RT-qPCR and qPCR during Las *in vitro* culture with and without the antibiotics. (D), Las growth inhibition measured by qPCR only during the cultivation. A significant difference between qRCR and RT-qPCR is indicated by different letters.

**Figure 7.**
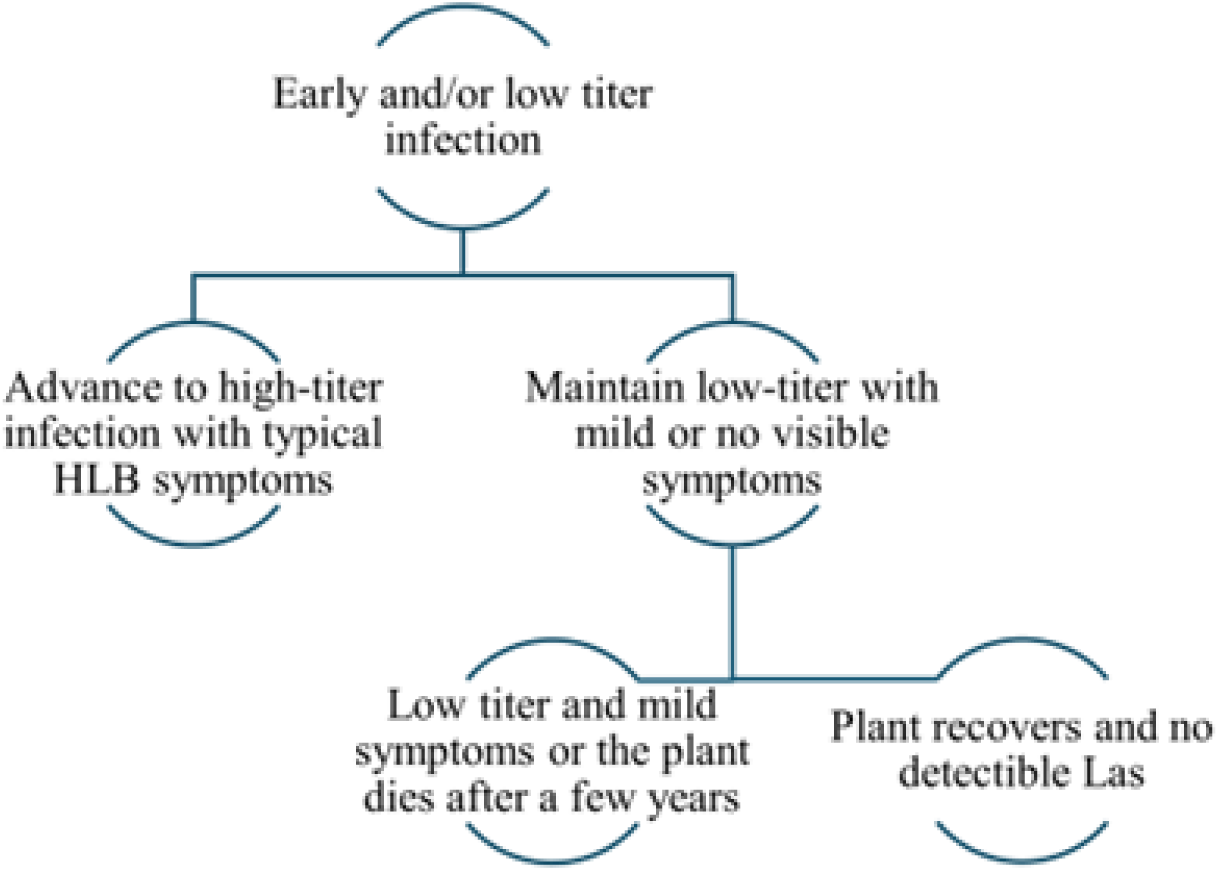
Schematic diagram of Las infection and progression in citrus plants.

## Discussion

### Improved sensitivity and specificity of the assay allows detection of the “undetectable”

RT-qPCR has the demonstrated capability to detect Las more sensitively and, in the face of the compounding effects of HLB on the citrus industry, this capability is especially important for early detection and detection of low-titer infections. The Li et al. (2006) detection method has been the preeminent detection method for over a decade despite the reported sensitivity of only 20 ng of total rDNA (Li et al. 2006), and the absence of a G nucleotide in the published primer sequence (Duan et al. 2009; Morgan et al. 2012). Different strategies have been developed in recent years attempting to improve the method including simple sampling strategy advances (Lin et al. 2010), utilization of single-tube nested qPCR (Lin et al. 2010), improved primer sets for qPCR based on new genetic information (Ananthakrishnan et al. 2013; Bao et al. 2020; Morgan et al. 2012; Orce et al. 2015; Park et al. 2018; Zheng et al. 2016), digital droplet qPCR (ddPCR) with detection of either one (Zhong et al. 2018) or two or more copies of targeted gene (Maheshwari et al. 2021; Selvaraj et al. 2018), and a CRISPR/Cas12a-based DETECTR assay (Wheatley et al. 2021). However, while sampling strategies should be optimized, they alone cannot improve the sensitivity of diagnostic assays. Sensitivity may be increased by using different primer sets in PCR-based systems, though extensive testing is required to validate new primers for diagnostic testing to ensure detection of all strains (Kunta et al. 2014b). While ddPCR is a promising new technology, its lower dynamic range renders ddPCR less suitable for high-throughput diagnostic testing than qPCR-based approaches. CRISPR/Cas detection mechanisms were first proposed in 2016 (Pardee et al. 2016; Yuan et al. 2022) and have yet to be widely adopted due to relatively inefficient original sample processing, relatively complex nucleic acid extraction requirements, the inability to detect multiple pathogens at once, and concerns over contamination in the Cas protein effect stage (Yuan et al. 2022). To overcome these challenges, RT-qPCR with existing validated primers may be a powerful tool. We have modified a detection system capable of detecting Las nucleic acids in as little as 10 picograms of TNA, even in high-background host DNA samples, and demonstrated its usefulness in detecting low-titer infections both early in the pre-symptomatic infection phase and for longer periods of latency and furthermore providing a measure of cellular activity.

For this analysis, we use the Ct value as an approximation of assay sensitivity as lower Ct values generally produce more reliable detection of lower titers within samples. Notably, we were able use the same primers and probes to reliably detect Las in samples with RT-qPCR that were, apparently, false-negatives in qPCR. When qPCR did detect Las in a sample, the RT-qPCR detection in the same sample was often at a lower Ct value. In the most contrasting cases, this enhanced detection could improve detection more than 500-fold over existing qPCR methods. The improvement in detection is especially important in citrus samples, where the false-negative rate in qPCR as compared to RT-qPCR is 18.40%. Many trees were tested at 49 dpi, prior to the onset of symptoms. Of these, the qPCR was able to detect Las in a small portion (3%). When tested with RT-qPCR, the early infections were more abundantly detected (29%). This trend remained constant for trees that had been diseased for longer, with greater or fewer HLB symptoms, and from several genetic backgrounds. While our results do indicate that qPCR with the Li 16S primers detected Las in most field-collected psyllid samples regardless of titer, the sensitivity of the assay remains superior for RT-qPCR over qPCR with RT-qPCR detecting Las in psyllids at an average of 5.74 cycles earlier than qPCR.

In the field, Las is spread by citrus psyllids as they migrate and feed. Psyllids are quick to feed and spread. Therefore, control up to this point has largely focused on eradicating psyllids and rogueing symptomatic trees. In some citrus producing regions, such as California in the US and São Paulo and Triângulo/Sudoeste Mineiro in Brazil, this has been the main method of HLB mitigation (Bassanezi et al. 2020). Previous research indicated that psyllids may transmit Las from trees that have yet to show symptoms (Lee et al. 2015), as well as from trees that provide only a low titer of Las in the phloem. In this case, removing only symptomatic trees in the absence of reliable psyllid control leaves many trees available as sources of inoculum for transmission of Las. Not only is the assay we developed sensitive enough to identify those trees that otherwise fall outside the range of detectability and thereby reduce false-negative detection, but it also could be combined with higher-throughput crude extraction methods (data not shown) yielding potentially lower concentrations of Las to make more widespread sampling feasible. The increased sensitivity additionally eliminates the need to excise and mince leaf midribs for the modified CTAB Las extraction, increasing the sample capacity. Midribs and laminar tissue differ by only a negligible ∼1 cycle in RT-qPCR (data not shown). Between the reduced time requirements involved in using laminar tissue, the high-throughput nature of RT-qPCR, and the ability of the method to detect smaller quantities of Las nucleic acids in leaf tissue, one-step RT-qPCR significantly reduces the amount of sampling, labor, and time required for accurate detection.

Accurate detection relies heavily on the familiarity of the collector with HLB symptoms, as HLB symptoms are similar to nutrient deficiencies such as zinc deficiency (Rao et al. 2018) which can be widespread in citrus-producing regions including Florida (Cuesta et al. 1993). In this study, experienced HLB researchers identified, categorized, and collected asymptomatic and symptomatic leaves. Meanwhile, leaves in the “general” category were collected by less-experienced individuals unfamiliar with HLB symptoms, thus reproducing more general sampling outcome. Las accumulates more heavily in symptomatic leaves, and the higher titers coupled with expert symptom identification allow for effective use of qPCR. Indeed, 100% of symptomatic leaves had detectable Las in both RT-qPCR and 16S rDNA-targeted qPCR. This represents a negligible error rate for true symptomatic tissue, but RT-qPCR becomes necessary in detection of samples taken from mixed collections of symptomatic and nutrient-deficient leaves. Our results indicate that qPCR of mixed sample collections may result in false negatives in approximately 15.45% of samples.

### Low-titer infections follow disparate progression patterns

In our work, several modes of HLB progression were observed. Based on our results, and as presented in **Error! Reference source not found.**, three to four outcomes seem possible after an initial low-titer Las infection. The first case describes the dogmatic transitory low-titer infection in which Las initially establishes itself at low titers as it moves through the phloem and replicates relatively fast to establish high titers and the onset of HLB symptoms within a few weeks. More novel is the description of persistent low-titer infections with mild to no symptoms, sometimes with apparent plant recovery. In trees inoculated with low-titer inoculum, low-titer infections could be maintained without significant increase for more than four years in an insect-free greenhouse. It has been previously demonstrated that pollen can carry active Las and transmit it without disease development (Wang et al. 2022), and that seeds harbor Las in all parts of the seed without resulting seedlings developing disease (Albrecht and Bowman 2009; Bagio et al. 2020). Additionally, to date Las has only been successfully grown in co-culture systems (Zheng et al. 2024). It is therefore possible that an unmixed infection by Las alone in citrus plants is insufficient to cause typical HLB, and that persistent low-titer infections were a result of Las colonization in the absence of co-infection with a helper bacterium.

### Assay improvement is dependent on relative 16S rRNA abundance

The greater sensitivity of RT-qPCR may rely on the abundance of RNA available for template. To test this, we looked at the performance of different primer sets targeting different genes. In direct comparisons of the same samples between the Li primers for 16S rRNA/rDNA, RNR primers, and LJ-900 primers, the Li primers and probe outperformed the other two, despite the higher copy number of genes in RNR targets (Maheshwari et al. 2021) and the multiple tandem repeats in the prophage genes *LasA_I_* and *LasA_II_* targeted by LJ-900 primers and probe (Morgan et al. 2012). Not only is the overall detection rate improved from 9% to 100% in low-titer citrus samples and 20% to 100% in low-titer psyllids, but the certainty of positives is also improved by the return of a lower Ct value. Certain samples, such as citrus leaves with a low titer of Las and particularly low titers associated with pre-symptomatic infection, may require the sensitivity of the Li RT-qPCR assay for detection. The Li primers are designed to target the 16S rDNA subunit of the Las ribosome, which is complementary to the Las 16S rRNA. During the reverse-transcription step in the one-step RT-qPCR reaction, the 16S rRNA transcripts are reverse-transcribed to cDNA. This would create a much more abundant source of Las DNA thereby artificially “increasing the copy number” to a much more detectable level.

### Cellular activity is linked to rRNA abundance as predicted by RT-qPCR/qPCR Ct gaps

Given that the increased detectability is related to the amount of rRNA present in a sample, it stands to reason that more active cells which necessarily have more ribosomes would have greater quantities of rRNA (Gourse et al. 1996; Pagliai et al. 2014) available for reverse-transcription than less active or inactive cells. Using this logic, the gap between the RT-qPCR measure of total nucleic acid titer and the qPCR measure of DNA titer can be used to develop an understanding of Las relative cellular activity in different hosts and potentially even tissues, timeframes, and population groups. Given the ease of sampling, extraction, and testing, this may provide a new system to test antimicrobial compounds outside of axenic culture for their effects on reducing cellular activity. Inactivation of Las cells *en masse* would likely result in more successful control of bacterial proliferation and spread. The mechanisms for Las interaction with its own population at different titers remain to be explored, but current evidence suggests that Las is most active in pre-symptomatic tissue (Folimonova and Achor 2010) as compared to symptomatic tissue. Research has shown that higher titers of Las are correlated with degeneration of the phloem system in a feedback loop that may limit Las mobility and activity. Our results demonstrate that there is likely a higher relative abundance of 16S rRNA in pre-symptomatic tissues than in symptomatic tissues, and that symptomatic tissues have similar 16S rRNA abundances regardless of the age of the leaves and symptomology.

In this study, we determined that relative Las cellular activity is likely higher in psyllids than in citrus as determined by Ct gaps. Across all 297 psyllid samples tested, the average Ct gap was 5.74 ±1.25 cycles. Across all citrus samples tested, the Ct gap was only 4.97 ±1.45 cycles. Psyllids are likely to return Ct values up to ∼3.5 cycles lower than citrus samples, indicating heightened Las cellular activity. This may be partially explained by Las action *in planta* or the different nutritional provisions of different microbiomes that may support different Las growth rates and pathogenicity. For example, while citrus phloem and psyllid hemolymph are similar in their chemical composition, the psyllid microbiome is high in trehalose and fatty acids whereas the citrus phloem microbiome is largely lacking these molecules (Killiny et al. 2016). These differences may promote greater cellular activity of Las in the psyllid hosts. In the Killiny et al. study, psyllid hemolymph composition was contrasted with that of the citrus phloem and the authors noted that these more plentiful nutrients in psyllids were a major difference between hemolymph and phloem. Previous attempts to culture Las have relied on extractions of seeds (Parker et al. 2014), phloem extracted from shoots (Davis et al. 2008), leaves (Sechler et al. 2009), and stem extracts (Ha et al. 2019), and more recently, psyllids (Molki et al. 2020). Although none of these methods have been widely reproducible, one of the keys to culture may be selecting the most active Las cells as the inoculum source. More research will need to be done to determine the mechanisms behind the improved Las activity in psyllids over citrus, however our data supports the use of psyllids as a Las culture inoculum source, and further testing with psyllid inocula is recommended.

During this study, many hundreds of samples have been examined and RT-qPCR has outperformed qPCR in each one. The reliability of the assay is demonstrated, and the application of the assay can be expanded to different aspects of Liberibacter-related research. Elucidation of the direct relationship between Ct gaps, cellular activity, and cellular viability will aid in the descriptive study of the Las lifecycle and the best methods to intervene with mitigation strategies. The adoption of the RT-qPCR method on a broad scale is recommended given the wide availability of the materials, including primers, probes, and manufactured master mix. The data presented from this study suggests that one-step RT-qPCR will be a vital tool in the control, mitigation, and potential eradication of Las. The ability to detect low-titer Las infections more simply, whether early or later in the infection process, provides the citrus-growing community with an important tool to study sources of inoculum in the field and provides the research community with additional resources for evaluating Las cultures and control measures.

## Abbreviations

ACP: Asian citrus psyllid
ANOVA: Analysis of Variance
CTAB: Cetyltrimethylammonium Bromide
ddPCR: digital droplet PCR
dpi: days post-inoculation
EDTA: Ethylenediaminetetraacetic acid
HLB: Huanglongbing
HSD: Tukey’s honestly significant difference
Las: Candidatus Liberibacter asiaticus
PCR: polymerase chain reaction
qPCR: quantitative PCR
RNR: ribonucleotide reductase
RT-qPCR: reverse transcription-quantitative PCR
SDS: sodium dodecyl sulfate
TNA: total nucleic acids
vPCR: viability PCR

## Declarations

### Ethics approval and consent to participate

Not applicable.

### Consent for publication

Not applicable.

### Authors’ Contributions

YD conceived and designed this research, RP, DZ, WL performed experiments and analyzed data, and all authors participated in the manuscript writing and have agreed to the published version of the manuscript.

## Acknowledgements

We are thankful to Spencer Marshall and Jane Zheng for their many technical contributions to the work.

---

**Supplementary Table 1.**
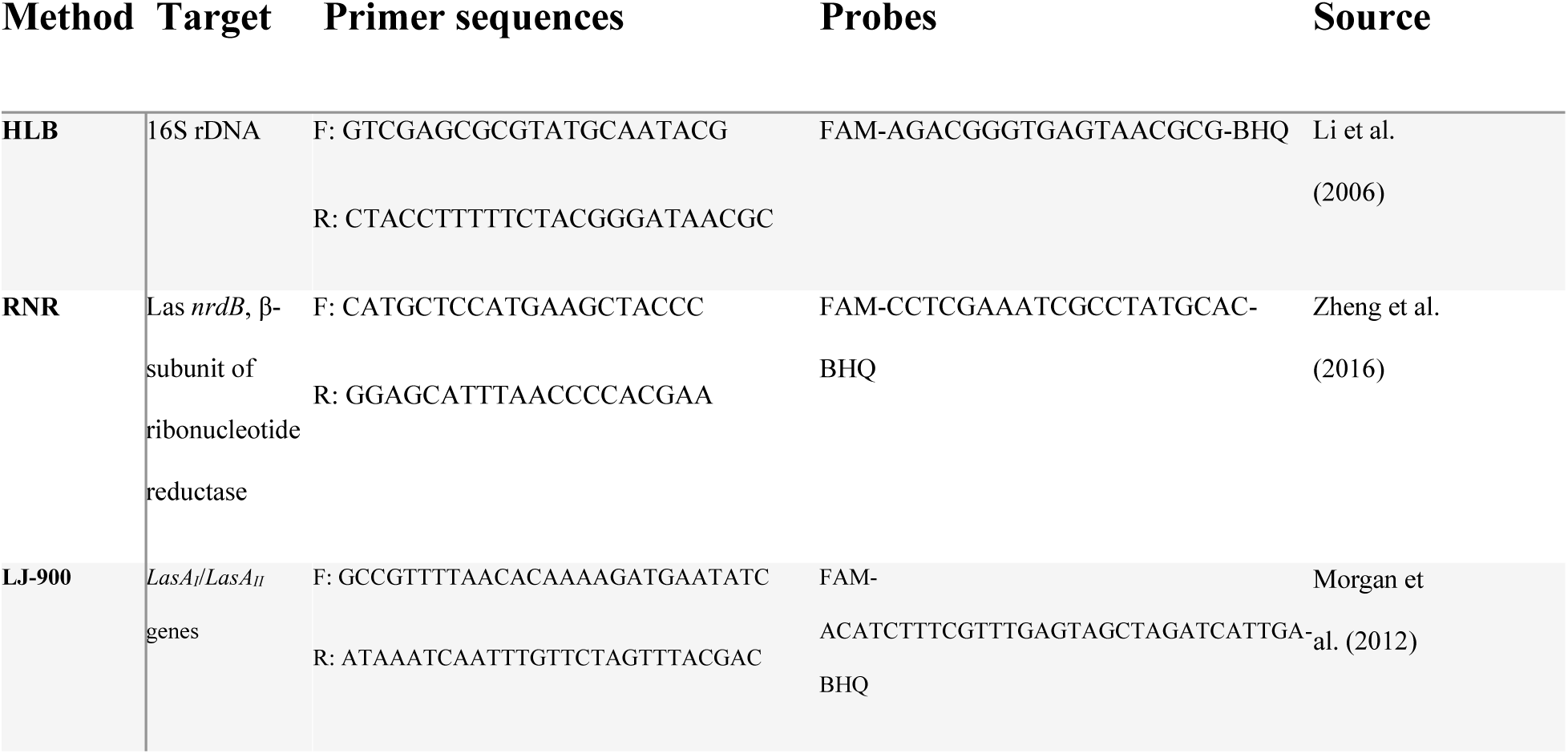

